# PRMT5 is a therapeutic target in choroidal neovascularization

**DOI:** 10.1101/2022.03.30.486275

**Authors:** Anbukkarasi Muniyandi, Matthew Martin, Mengyao Sun, Kamakshi Sishtla, Nathan R Jensen, Aishat Motolani, Xiaoping Qi, Michael E Boulton, Lakshmi Prabhu, Tao Lu, Timothy W Corson

## Abstract

Ocular neovascular diseases including neovascular age-related macular degeneration (nvAMD) are widespread causes of blindness. Patients’ non-responsiveness to currently used biologics that target vascular endothelial growth factor (VEGF) poses an unmet need for novel therapies. Here, we identify protein arginine methyltransferase 5 (PRMT5) as a novel therapeutic target for nvAMD. PRMT5 is a well-known epigenetic enzyme. We previously showed that PRMT5 methylates and activates a proangiogenic and proinflammatory transcription factor, the nuclear factor kappa B (NF-κB), which has a master role in tumor progression, notably in pancreatic ductal adenocarcinoma and colorectal cancer. We identified a potent and specific small molecule inhibitor of PRMT5, PR5-LL-CM01, that dampens the methylation and activation of NF-κB. Here for the first time, we assessed the antiangiogenic activity of PR5-LL-CM01 in ocular cells. Immunostaining of human nvAMD sections revealed that PRMT5 is highly expressed in the retinal pigment epithelium (RPE)/choroid where neovascularization occurs, while mouse eyes with laser induced choroidal neovascularization (L-CNV) showed PRMT5 is overexpressed in the retinal ganglion cell layer and in the RPE/choroid. Importantly, inhibition of PRMT5 by PR5-LL-CM01 or shRNA knockdown of PRMT5 in human retinal endothelial cells (HRECs) and induced pluripotent stem cell (iPSC)-derived choroidal endothelial cells (iCEC2) reduced NF-κB activity and the expression of its target genes, such as tumor necrosis factor α (TNF-α) and VEGF-A. In addition to inhibiting angiogenic properties of proliferation and tube formation, PR5-LL-CM01 blocked cell cycle progression at G1/S-phase in a dose-dependent manner in these cells. Thus, we provide the first evidence that inhibition of PRMT5 impedes angiogenesis in ocular endothelial cells, suggesting PRMT5 as a potential therapeutic target to ameliorate ocular neovascularization.

## 1 Introduction

Neovascular age-related macular degeneration (nvAMD) is a major cause of blindness, affecting an estimated 20 million older adults worldwide. It is characterized by macular neovascularization (MNV), the abnormal growth of new blood vessels (angiogenesis) from the choriocapillaris into the sub-retinal pigment epithelium (RPE) and/or subretinal space. nvAMD causes progressive degeneration of the neuroretina and RPE-choroid complex, leading to central vision loss (Keenan et al., 2021). Angiogenesis and inflammation are the hallmarks of nvAMD, where angiogenesis triggers endothelial cell proliferation, migration, and vasculogenesis along with dysregulation of vascular endothelial growth factor (VEGF) as a key pathological factor (Kinnunen and Ylä-Herttuala, 2012). Ocular neovascularization also underlies other blinding eye diseases, such as proliferative diabetic retinopathy (PDR) and retinopathy of prematurity (ROP) (Campochiaro, 2015). Although a number of antiangiogenic therapies targeting the VEGF signaling cascade yield successes in many patients, the anti-VEGF biologics still have significant shortcomings due to non-responsive patients and drugs’ tachyphylaxis (Nagai et al., 2016). Therefore, there remains a critical need to uncover novel therapeutic targets for better understanding and treating pathological ocular angiogenesis. Inflammatory signaling represents an appealing area in which to discover such therapeutic targets.

The nuclear factor (NF)-κB constitutes a family of inducible transcription factors that mediate the expression of a variety of genes involved in inflammation, cell growth and development, angiogenesis, and other pathways (Motolani et al., 2021). The five NF-κB family members, which include RelA/p65, RelB, c-Rel, p50/p105 (NF-κB1), and p52/p100 (NF-κB2), share the same REL homology domain (RHD) that is responsible for DNA binding and dimerization (Hartley et al., 2020). These transcription factors dimerize to regulate gene expression in two distinct NF-κB signaling pathways: the canonical pathway and the non-canonical pathway (Martin et al., 2021). The canonical pathway, characterized by its quick and transient activity, is activated by cytokines, growth factors, and ligands of pattern recognition receptors (PRRs), TNF receptor (TNFR) superfamily members, T-cell receptor (TCR), and B-cell receptor (Liu et al., 2017). Following the activation of the canonical pathway, a series of phosphorylation cascades involving IκB kinase (IKK) and the inhibitor of nuclear factor kappa B alpha (IκBα) leads to the translocation of p65/p50 into the nucleus, where they bind the promoters of their target genes and activate gene transcription (Liu et al., 2017). On the other hand, the activation of the non-canonical pathway is mediated by the RelB/p52 complex and occurs in response to stimuli from ligands of receptors such as lymphotoxin β receptor (LTβR), BAFF receptor (BAFFR), CD40 and receptor activator of NF-κB (RANK). Thus, this pathway is highly involved in biological functions such as lymphoid organ development, B-cell survival and maturation, and bone metabolism (Sun, 2011).

Multiple reports have demonstrated the extensive role of NF-κB, particularly the prototypical subunit p65, in the pathogenesis of different diseases. Given its critical role in inflammation, NF-κB contributes to the development and/or the progression of conditions such as asthma, arthritis, inflammatory bowel diseases, atherosclerosis, Alzheimer’s disease, cancer, and diabetes. In some cancers, the constitutive activation of NF-κB correlates with poor clinical course and outcomes (Yamamoto and Gaynor, 2001). Similarly, NF-κB activation controls the expression of proangiogenic genes, whose aberrant expression underlies the pathology of certain eye diseases (Stoltz et al., 1996; Xie et al., 2010; Lan et al., 2012). Many studies have investigated the association of nvAMD with NF-κB in clinical samples and in AMD experimental models *in vitro* and *in vivo*. They show that activation of NF-κB induced inflammatory signaling and upregulated tumor necrosis factor α (TNF-α) and VEGF in retinal pigment epithelial (RPE) cells (Oh et al., 1999; Cousins et al., 2004; Shi et al., 2006; Liu et al., 2014; Lu et al., 2014; Wang et al., 2015; Wang et al., 2016; Ghosh et al., 2017; Wang et al., 2017; Hikage et al., 2021; Sharma et al., 2021; Xin et al., 2021). Thus, exploring key therapeutic druggable targets that directly mediate NF-κB regulation in AMD pathology, and their potent small molecule inhibitors would be highly valuable in the discovery of new therapeutics targeting nvAMD.

Previously, we identified protein arginine methyltransferase 5 (PRMT5) as a novel activator of NF-κB in cancer (Wei et al., 2014; Prabhu et al., 2017b). We speculate that PRMT5 could also play an important role in ocular neovascularization through its regulation of NF-κB. A type II arginine methyltransferase, PRMT5 belongs to the PRMT superfamily, initially named as Janus-kinase binding protein (JBP1) (Wei et al., 2014). PRMT5 is a multifunctional protein well-known for methylating histones to regulate their transcriptional activities. To date, PRMT5 has been shown to play a significant role in tumorigenesis and is overexpressed in multiple types of cancers including colon (Prabhu et al., 2017b; Yan et al., 2021), lung (Li et al., 2019), liver (Jiang et al., 2018), pancreas (Prabhu et al., 2017b; Qin et al., 2019), kidney (Zhang et al., 2018), brain (Han et al., 2014), and others. However, the role of PRMT5 in ocular neovascular diseases has never been examined before. PRMT5 is also known to have a critical role in the dimethylation of p65 (*i.e.*, p65me2) and plays a role in hyperactivation of NF-κB, promoting NF-κB mediated cell signaling pathways (Wei et al., 2013). Given this proinflammatory role, we expect the activity of PRMT5 on NF-κB activation to be increased in the setting of ocular neovascularization. Thus, targeting PRMT5 by novel inhibitors might render a beneficial therapeutic effect.

We developed a high throughput AlphaLISA screen approach and identified a potent small molecule inhibitor of PRMT5, PR5-LL-CM01. We showed that PR5-LL-CM01 reduced pancreatic and colorectal tumor phenotypes by inhibiting the activation of NF-κB and its downstream targets (Prabhu et al., 2017a; Prabhu et al., 2017b). PR5-LL-CM01 demonstrated binding affinity to the active site of PRMT5 *in silico*, and it displayed anti-tumor effects in both *in vitro* and *in vivo* systems of pancreatic ductal adenocarcinoma (PDAC) and colorectal cancer (CRC) superior to the commercial inhibitor of PRMT5, EPZ015666 (Prabhu et al., 2017b). We therefore propose PR5-LL-CM01 as a potential means to inhibit ocular neovascularization.

In this study, we show that PRMT5 is highly expressed in human and murine choroidal neovascularization and its genetic or chemical inhibition reduces proliferation, tube formation, NF-κB activation, and the expression of its target genes in ocular endothelial cells including human retinal microvascular endothelial cells (HRECs) and induced pluripotent stem-cell (iPSC) derived choroidal endothelial cells (iCEC2). Moreover, PR5-LL-CM01 blocked cell cycle progression at G1/S-phase in a dose-dependent manner in these cells. Collectively, our data suggest that PR5-LL-CM01 can be explored as a novel treatment lead for targeting PRMT5-mediated inflammation and angiogenesis in ocular neovascular diseases.

## 2 Materials and Methods

### 2.1 Animals

All animal studies were approved by the Institutional Animal Care and Use Committee, Indiana University School of Medicine and the “Use of Animals in Ophthalmic and Visual Research” guidelines of the Association for Research in Vision and Ophthalmology were followed. Wild-type C57BL/6J mice (female, 7 weeks of age) were purchased from Jackson Laboratory (Bar Harbor, ME, USA) and housed under standard conditions (Wenzel et al., 2015) at the Laboratory Animal Resource Center, Indiana University School of Medicine.

### 2.2 Laser-induced choroidal neovascularization (L-CNV) model

L-CNV was done as described before (Basavarajappa et al., 2017; Sardar Pasha et al., 2018) with minor modifications in laser power and duration. Briefly, mice were anesthetized by intraperitoneal injections of ketamine hydrochloride (80 mg/kg) and xylazine (10 mg/kg). The pupils of the eyes were dilated using tropicamide (1%) and phenylephrine (2.5%) and the eyes were exposed to laser treatment with 270 mW power pulses of the Micron IV laser injector (Phoenix Research Labs, Pleasanton, CA, USA) using 532 nm wavelength, 70 ms duration and 50 μm spot size. Mice were euthanized and eyes removed at day 1, 7 and 14. The untouched control mice underwent enucleation at the end of the experiment on day 14. Eyes were fixed in 4% paraformaldehyde (PFA).

### 2.3 Flat-mount staining

The RPE/choroids and retinas were dissected from the fixed eyes and prepared as flat mounts for PRMT5 and isolectin B4 (IB4) vasculature staining. In brief, the retina and choroid were fixed again with 4% PFA overnight at 4°C. The tissues were then washed with phosphate-buffered saline (PBS, 1X) twice for 10 minutes each, permeabilized in blocking buffer containing 5% bovine serum albumin (BSA) and 0.5% Triton X-100 in PBS for 70 minutes. After blocking, the tissues were stained with isolectin B4 from *Griffonia simplicifolia* (GS-IB4; Biotin-conjugated, 1:250 dilution, Invitrogen, Waltham, MA, USA #121414) and anti-PRMT5 (1:150 dilution, Abcam, Waltham, MA, USA #ab109451; RRID:AB_10863428) prepared in antibody blocking solution containing 0.5% BSA (Fisher, Pittsburgh PA, USA #BP9703-100) and 0.5% Triton X-100 (Sigma-Aldrich, St. Louis, MO, USA #T9284) for 48-72 hours at 4°C in a shaker. Four times for 15 min each, the tissues were washed with 1× PBS (Thermo Scientific, Waltham, MA, USA #10010-023) and incubated overnight in a shaker protected from light at 4°C with respective detection reagents (DyLight 488-conjugated streptavidin at 1:400 dilution, Invitrogen #21832; and Alexafluor 555-conjugated goat anti-rabbit antibody at 1:250 dilution, Invitrogen #A21428; RRID:AB_2535849) prepared in antibody blocking solution. Then the tissues were washed with 1× PBS for four times, 15 minutes each and finally the immunostained tissues were flat mounted on glass slides and cover-slipped using Fluoromount-G (Southern Biotechnology, Birmingham, AL, USA). Until imaging, the flat mounts were protected from light and stored at 4°C. The immunostained L-CNV lesions were imaged by confocal microscope (LSM700, Carl Zeiss, Thornwood, NY, USA) using 20× objective and Z-stack optical sectioning.

### 2.4 Immunostaining

Deidentified human donor eyes from nvAMD patients and age-matched controls were sourced from the National Disease Research Interchange (NDRI, PA, USA), which obtains informed consent for donor materials following the principles stated in the WMA Declaration of Helsinki and the Department of Health and Human services Belmont Report. The sections of the eye were deparaffinized, rehydrated and underwent antigen unmasking by heat-induced epitope retrieval in 1× citrate buffer (pH 6.0; Thermo Scientific, Waltham, MA, USA #AP-9003-500). The sections were rinsed with PBS and blocked for 2 hours at room temperature using 10% normal donkey serum (NDS; Abcam, #ab7475) prepared in 1% BSA in PBS, then incubated with primary (rabbit anti-PRMT5 at 1:100 dilution; Abcam, Waltham, MA, USA #ab109451; RRID:AB_10863428) and rabbit IgG control (1:100 dilution; R&D systems, Minneapolis, MN, USA # AB-105-C; RRID:AB_354266) antibodies prepared in 10% NDS plus 1% BSA for overnight at 4°C. Sections were washed thrice with PBS-T (PBS+0.025% Tween-20) for 5 minutes each. They were then incubated with Alexafluor 555-conjugated goat anti-rabbit secondary antibody (Invitrogen) at 1:200 dilution, for 1 hour in a dark humidified chamber at room temperature, followed by washing thrice with PBS-T for 5 minutes each. Finally, the sections were dehydrated through an ethanol series and mounted with Vectashield mounting medium with the nuclear stain, DAPI.

The mouse eyes harvested after L-CNV induction at different days along with the eyes from untouched control mice were fixed in 4% PFA (Thermo Fisher Scientific, #43368) for 16 hours at 4°C. Dissected choroid and retina were embedded in optimal cutting temperature compound and cryosectioned to 5 μm thickness. The same immunostaining protocol described above was followed excluding deparaffinization, rehydration and antigen unmasking steps. The reagents used for mouse cryosections were as follows: primary rabbit anti-PRMT5/rabbit IgG control (1:150 dilution), GS-IB4 (1:250 dilution) and secondary Alexafluor 555-conjugated goat anti-rabbit antibody (1:250 dilution) and DyLight 488-conjugated streptavidin (1:400 dilution). The mounted human and mouse sections were imaged in a confocal microscope (LSM700, Carl Zeiss, Thornwood, NY, USA) with a 20× objective.

### 2.5 Immunoblotting for tissues

Immunoblotting from the mouse eye lysates was performed as described before (Pran Babu et al., 2020). Briefly, retina and choroid tissues dissected out from the L-CNV and untouched control mice (four eyes) were lysed for 20 min on ice in radioimmunoprecipitation assay (RIPA) buffer (Sigma #R0278) with protease inhibitors (cOmplete mini, #04693159001, Roche, Indianapolis, IN, USA) and phosphatase inhibitors (PhosSTOP, #04906837001, Roche). The samples were homogenized and centrifuged at 12,000 *g* for 15 min at 4°C. Supernatants (lysates) were separated, and protein concentrations were determined using bicinchoninic acid (BCA) protein assay. Equal amounts of protein (25-35 μg) from retina and choroid samples were resolved by 10% SDS-PAGE and transferred onto polyvinylidene fluoride (PVDF) membranes (Millipore, Burlington, MA, USA). The antibodies against PRMT5 (rabbit, 1:1000 dilution, Abcam, Waltham, MA, USA #ab109451) and β-actin (mouse, 1:1000 dilution, Sigma-Aldrich, #A5316; RRID:AB_476743) were used to immunoblot the proteins with secondary antibodies, anti-rabbit IgG peroxidase conjugated (1:10,000, Rockland, Rockland, MD, USA, #611-1302; RRID:AB_219720) and anti-mouse IgG peroxidase conjugated (1:10,000, Rockland, #610-1302; RRID:AB_219656). All of the antibody dilutions were prepared in Tris-buffered saline containing 2.5% BSA. Amersham ECL prime immunoblotting detection reagents (#RPN2236, GE Healthcare, Chicago, IL, USA) were used to detect immunoreactive bands on a ChemiDoc MP imaging system (Bio-Rad, Hercules, CA, USA).

### 2.6 Cell culture

The HRECs were purchased from Cell Systems (Kirkland, WA, USA) and grown in endothelial basal medium (EBM) in combination with endothelial cell growth medium (EGM-2) bullet kit (Lonza, Walkersville, MD, USA). iCEC2 cells were a kind gift from Dr. Robert F. Mullins’ laboratory (Voigt et al., 2019) at the University of Iowa and grown in endothelial cell growth medium (R&D Systems, Minneapolis, MN, USA). iCEC2 cells contain a temperature-sensitive hypomorphic T antigen and actively proliferate at 33°C, which was used for stock maintenance, while growth slows at 37°C, which was used for functional analyses. All cells were tested for mycoplasma contamination regularly.

### 2.7 Construction of stable cells

FLAG-tagged WT-PRMT5 cDNA or FLAG-tagged WT-p65 cDNA was amplified by reverse transcription from total mRNA derived from 293 cells (Wei et al., 2013). The sequences were confirmed via DNA sequencing and then cloned into respective pLVX-IRES-puro vectors. Lentiviral vectors expressing five different PRMT5-directed shRNAs (target set RHS4533-EG10419), and the universal negative control, pLKO.1 (RHS4080) were purchased from Open Biosystems (Dharmacon, Lafayette, CO, USA). To generate stable cells, the lentiviral plasmid containing the DNA of interest or shRNAs targeting PRMT5 exons were transfected into a 293T packaging cell line to produce viruses. HREC or iCEC2 cells were then infected with these viruses and further selected with 1 μg/ml of puromycin, as the lentiviral vector construct includes a puromycin resistance gene. Expression of the respective constructs was confirmed using immunoblotting with specific antibodies. Cells were used for further experiments, with HRECs used between passages 5 and 7.

### 2.8 Proliferation assays

HRECs and iCEC2 cells overexpressing PRMT5 and shPRMT5 knockdown cell lines were plated at 2 × 10^4^ cells/well in a 6-well plate. Cells were seeded in triplicate and counted on different days using a hemocytometer. For compound effects on proliferation, HRECs and iCEC2 cells were evaluated as described earlier (Basavarajappa et al., 2015; Basavarajappa et al., 2017; Sardar Pasha et al., 2018). In brief, the cells were seeded in 96-well black plates with clear bottom at a density of 2.5 × 10^3^ with 100 μl of growth media and incubated for a day. After 24 h, 1 μl of PR5-LL-CM01 (0.1 nM to 100 μM) or vehicle DMSO (at 1% final concentration to the cells) was added, and the plates were incubated for 44 h. To each well of the plates, 11.1 μl of Alamar blue reagent was added and the readings were taken after 4 hours in a Synergy H1 plate reader (BioTek, Winooski, VT) with 560 nm (excitation) and 590 nm (emission) wavelengths. Using GraphPad Prism 9.0, GI_50_ (growth inhibitory concentration) was calculated for PR5-LL-CM01.

### 2.9 Flow cytometry cell cycle analysis

The analysis of cell cycle in HRECs and iCEC2 cells was assessed as described before (Sardar Pasha et al., 2018). Briefly, the cells were grown in 6 well plates (2 × 10^6^) and at 70% confluency, the cells were treated with PR5-LL-CM01 (0.1 μM, 1 μM and 10 μM) or vehicle DMSO (1% final concentration to the cells) for 24 h. The cells were washed with PBS twice and fixed in 70% ethanol. Prior to analysis, the cells were washed with PBS for two times, the cell pellets resuspended in propidium iodide (PI) staining solution (20 μg/ml) prepared in 1× PBS containing 100 μg/ml of RNAse A and 0.1% Triton X-100) and incubated at room temperature for 30 min. The cells were then analyzed by Attune NxT flow cytometer (Thermo Fisher Scientific, Waltham, MA, USA). The single cell population was analyzed by area histograms and cell cycle profiles were created using Modfit software (v. 5.0, Verity Software House, Topsham, ME, USA). Pulse shape analysis was performed to eliminate any debris, doublets and aggregates from the whole cell population analyzed.

### 2.10 Matrigel tube formation assay

The tube formation ability of HRECs, iCEC2 cells and the respective stable cells was assessed as previously described (Basavarajappa et al., 2017; Sardar Pasha et al., 2018). Briefly, the wells of a 96 well plate were precoated with Matrigel (50 μl), which was allowed to solidify at 37°C for 20 min. The cells were seeded at a density of 1.5 × 10^4^ in growth media (100 μl) containing 1 μl/well PR5-LL-CM01 (0.1 μM, 1 μM and 10 μM) or vehicle DMSO (1% final concentration to the cells). The stable cells were seeded at the same density over Matrigel basement membrane. Tube formation was monitored every 2 hours, after incubating the cells for 8 h at 37°C and 5% CO_2_, each well was photographed using a brightfield microscope (4× objective), and the measurements of formed tubes were analyzed using Angiogenesisanalyzer plugin in ImageJ software (v. 1.8.0; http://image.bio.methods.free.fr/ImageJ/?Angiogenesis-Analyzer-for-ImageJ.html).

### 2.11 Immunoblotting for cells

HRECs and iCEC2 cells were pelleted post treatment with PR5-LL-CM01 (0, 1.5, 3, or 6 μM respectively for 24 hours) in phosphate buffered saline (PBS). Pellets were lysed with lysis buffer [(10mM Tris-Cl pH 8.0, 1 mM EDTA, 1% Triton X-100, 0.1% sodium deoxycholate, 0.1% SDS (sodium dodecyl sulfate), 14 mM NaCl, 1 mM phenylmethylsulfonyl fluoride]. Protein concentration for each sample was determined using Protein Assay Reagent (Bio-Rad, Hercules, CA, USA). Equal protein concentrations were run on a 10% SDS-PAGE (polyacrylamide gel electrophoresis) gel and transferred to a polyvinylidene difluoride membrane (Thermo Fisher Scientific). Membranes were exposed to anti-p65, anti-p65me2, anti-flag, anti-PRMT5, and anti-actin and their respective secondary antibodies in mouse or rabbit respectively. based on manufacturer’s instructions. Protein signal was detected using enhanced chemiluminescence (ECL) reagent (PerkinElmer). The antibodies used were obtained from commercial sources: Rabbit anti-p65 (at 1: 3,000 dilution; Santa-Cruz Biotech, Dallas, TX, USA, #SC-109), rabbit anti-p65me2 (customized antibody, Genscript, Piscataway, NJ), mouse anti-FLAG M2 (at 1:3,000 dilution; Millipore-Sigma, MO, USA, #F-1804), mouse anti-β-actin (at 1: 5,000 dilution; Sigma-Aldrich, St. Louis, MO, USA, #A5316), rabbit anti-PRMT5 (at 1:3,000 dilution; Abcam, Waltham, MA, USA #ab109451), goat anti-rabbit IgG (H+L) secondary antibody, HRP (at 1:3,000 dilution; ThermoFisher Scientific, Waltham, MA, USA, # 31460), goat anti-mouse IgG (H+L) secondary antibody, HRP (at 1:3,000 dilution; ThermoFisher Scientific, Waltham, MA, USA, # 62-6520).

### 2.12 NF-κB luciferase assay

The NF-κB luciferase lentiviral construct pLA-NFκBmCMV-luc-H4-puro (Lu et al., 2010) (containing five tandem copies of the NF-κB site from the IP10 gene) was introduced in the respective cells using Lipofectamine™ LTX Reagent and PLUS Reagents (Thermo Fisher Scientific). Luciferase activity was measured after 48 hours (with or without drug treatment) using Reporter Lysis Buffer kit (Promega, Madison, WI) per manufacturer’s instructions and a Synergy H4 plate reader.

### 2.13 Quantitative reverse-transcription polymerase chain reaction (qRT-PCR)

Following treatment with PR5-LL-CM01 for 24 hours, total RNA was isolated from the respective cells using TRIzol. First strand complementary DNA was generated using SuperScript III First-Strand Synthesis Kit (Invitrogen, Carlsbad, CA). GAPDH was selected as the housekeeping gene for normalization; each gene was run along with GAPDH, and the difference between threshold cycles (CT) was designated as ΔC_T_. ΔΔC_T_ is the difference between their respective controls. qPCR was subsequently executed using FastStart Universal SYBR Green kit (Roche, Indianapolis, IN). Primers were designed using the Primer Express 3.0 software (Thermo Fisher Scientific). GAPDH-F CCATCACCATCTTCCAGGAGCG; GAPDH-R AGAGATGATGACCCTTTTGGC; VEGFA-F TTGCCTTGCTGCTCTACCTCCA; VEGFA-R GATGGCAGTAGCTGCGCTGATA; TNFA-F TGGCCCAGGCAGTCAGA; TNFA-R GGTTTGCTACAACATGGGCTACA.

### 2.14 Statistical analyses

The data were analyzed using GraphPad Prism software (version 9.2.0). Unpaired Student’s *t*-test with Welch’s correction was used for comparing two means, while one-way ANOVA with Dunnett’s post hoc tests was used for comparing more than two means. Two-sided p values (p<0.05) were considered statistically significant.

## 3 Results

### 3.1 PRMT5 is highly expressed in nvAMD

To investigate the potential functional role of PRMT5 in ocular neovascularization, we first sought to examine its expression in postmortem eyes from human nvAMD patients. Sections of human nvAMD eyes in comparison with healthy control eyes showed PRMT5 expression in all the layers of the retina, with especially high expression seen in the RPE-choroid where neovascularization originates in nvAMD (Figure 1A and B). Also, AMD eyes compared to healthy control eyes clearly display the distinctive degenerative phenotypes, including the loss of inner of outer segments of photoreceptors and disruption in the retinal nuclear layer architecture (Figure 1A), although these changes and the pattern of PRMT5 expression vary among the AMD patients we studied.

**Figure 1.**
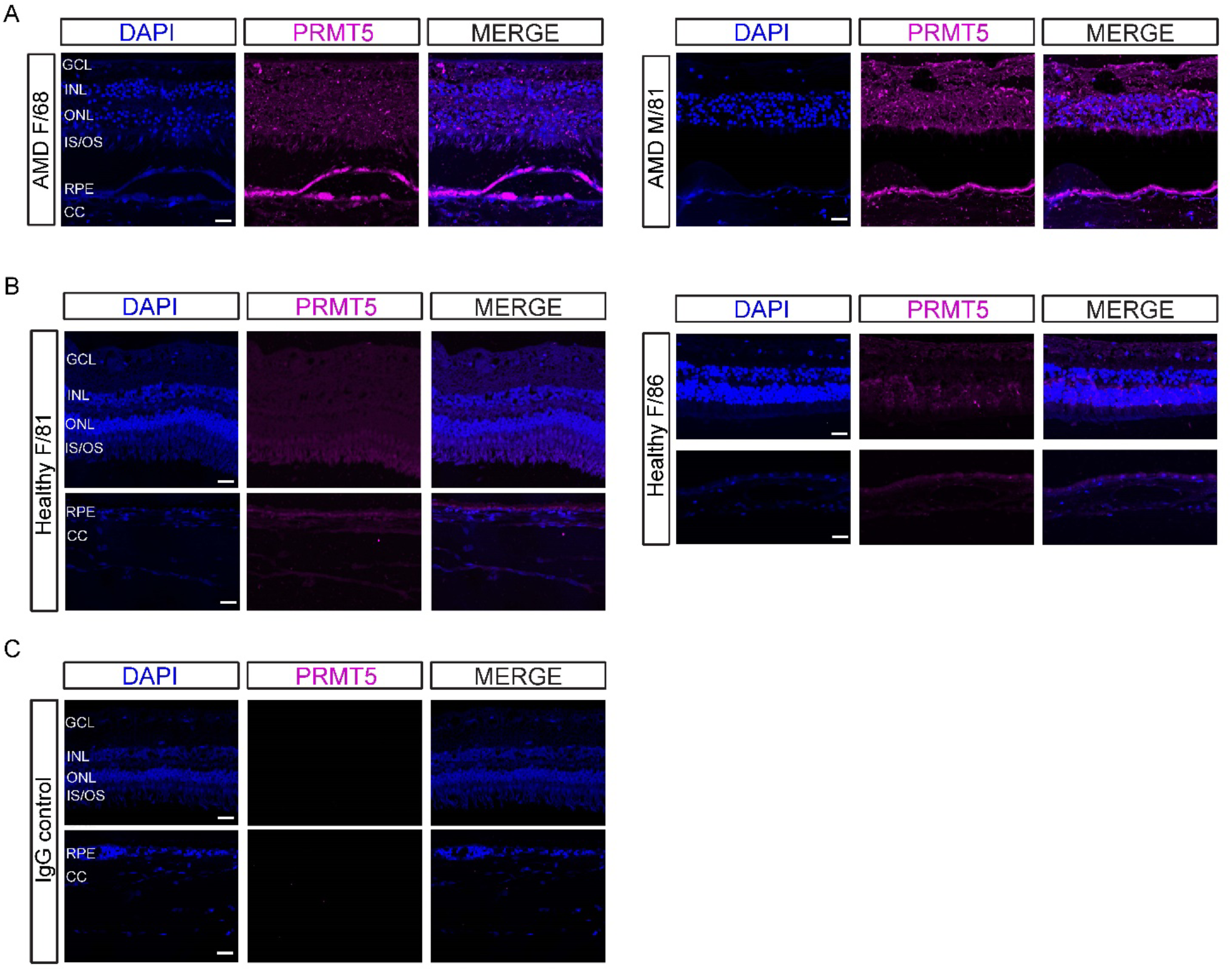
PRMT5 is highly expressed in neovascular age-related macular degeneration (nvAMD). PRMT5 immunostaining on sections of eyes from human (A) nvAMD patients (68 year old female and 81 year old male) and (B) age-matched controls (81 year old female and 86 year old female), where DAPI (blue) shows the nuclei of the cells and magenta indicates PRMT5 expression in different layers of the retina, and in the RPE/choroid complex. Higher expression of PRMT5 is observed in the retinal pigment epithelium (RPE)/choroid in the eyes with nvAMD. Representative samples shown from n=3 nvAMD patients and age-matched controls. (C) shows preimmune IgG control. Scale bar = 20 μm. GCL, ganglion cell layer; INL, inner nuclear layer; ONL, outer nuclear layer; IS/OS photoreceptor inner/outer segments; CC, choriocapillaris.

### 3.2 PRMT5 is highly expressed in murine L-CNV

Given the expression of PRMT5 in nvAMD, we next asked whether PRMT5 was similarly upregulated in experimental CNV. We studied this using a mouse model of L-CNV, wherein the mice eyes were harvested at 7 days post laser treatment and assessed for the expression of PRMT5 in the retina and choroid. Overexpression of PRMT5 was observed in and around the L-CNV lesions in choroidal flat mounts compared to the flat mounts prepared from untouched control eyes (Figure 2A). Immunoblot in comparison with untouched control eyes showed relatively increased levels of PRMT5 both in retina and choroid (Figure 2B) in the L-CNV samples. Immunostaining of PRMT5 on mouse eye cryosections revealed high expression and localization of PRMT5 in the ganglion cell layer (GCL) of the retina and in the RPE-Bruch’s membrane (BM)-choroid complex in L-CNV, although some expression was seen throughout the retina (Figure 2C).

**Figure 2.**
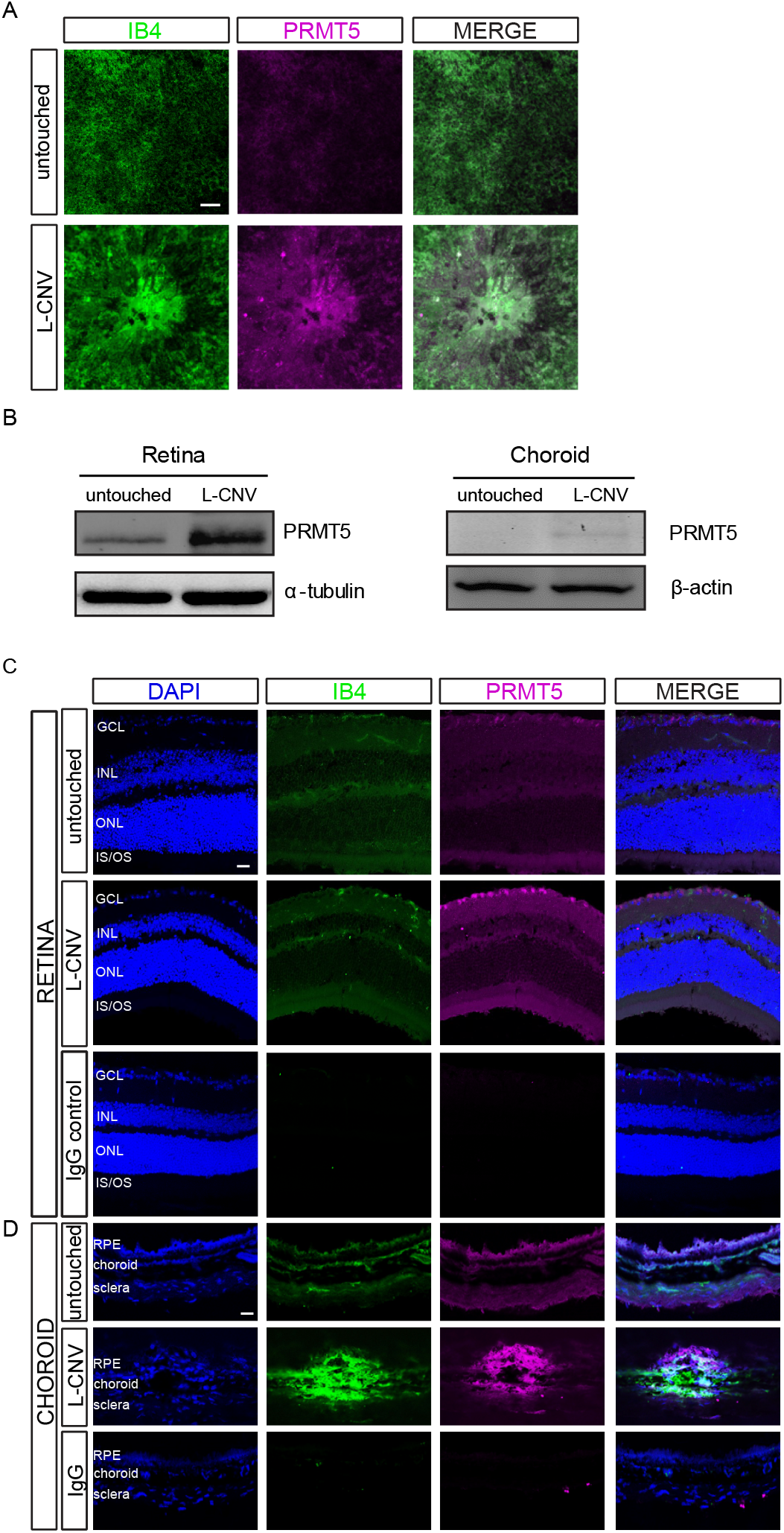
PRMT5 expression in murine laser-induced choroidal neovascularization (L-CNV). (A) Flat mount staining of L-CNV choroids, showing the expression of PRMT5 (magenta) in and around the neovascular lesion in the choroid that underwent laser treatment compared to untouched control. (B) Immunoblot, showing PRMT5 is highly expressed in the retina and choroid of the L-CNV mouse eyes compared to the untouched control eyes. (C, D) Cryosections of the L-CNV (C) retina and (D) choroid, showing PRMT5 expression in different layers of the retina, including the ganglion cell layer (GCL), inner plexiform layer (IPL), outer plexiform layer (OPL), and in the inner and outer segments (IS/OS) of photoreceptors, and in the retinal pigment epithelium (RPE)-choroid complex where neovascularization is observed (isolectin B4 [IB4] staining, green). Higher expression is observed in the GCL and in the RPE-choroid in L-CNV compared to untouched.

### 3.3 PRMT5 regulates endothelial cell proliferation

Due to the expression profile of PRMT5 in CNV we generated PRMT5 overexpression and shPRMT5 knockdown in HRECs and iCEC2 cells. We were able to successfully overexpress a FLAG-tagged PRMT5 protein in HREC and iCEC2 cells compared to a vector control and to knock down PRMT5 compared to shScramble controls (Figure 3A and B). These cells were then used in future experiments. As NF-κB is an important regulator for cell growth, we first performed cellular proliferation assays (Figure 3C-F). Overall, overexpression of PRMT5 increased cellular growth compared to the vector control, while shRNA knockdown of PRMT5 decreased growth compared to the shScramble control. These data suggest PRMT5 plays an important role in promoting ocular endothelial proliferation.

**Figure 3.**
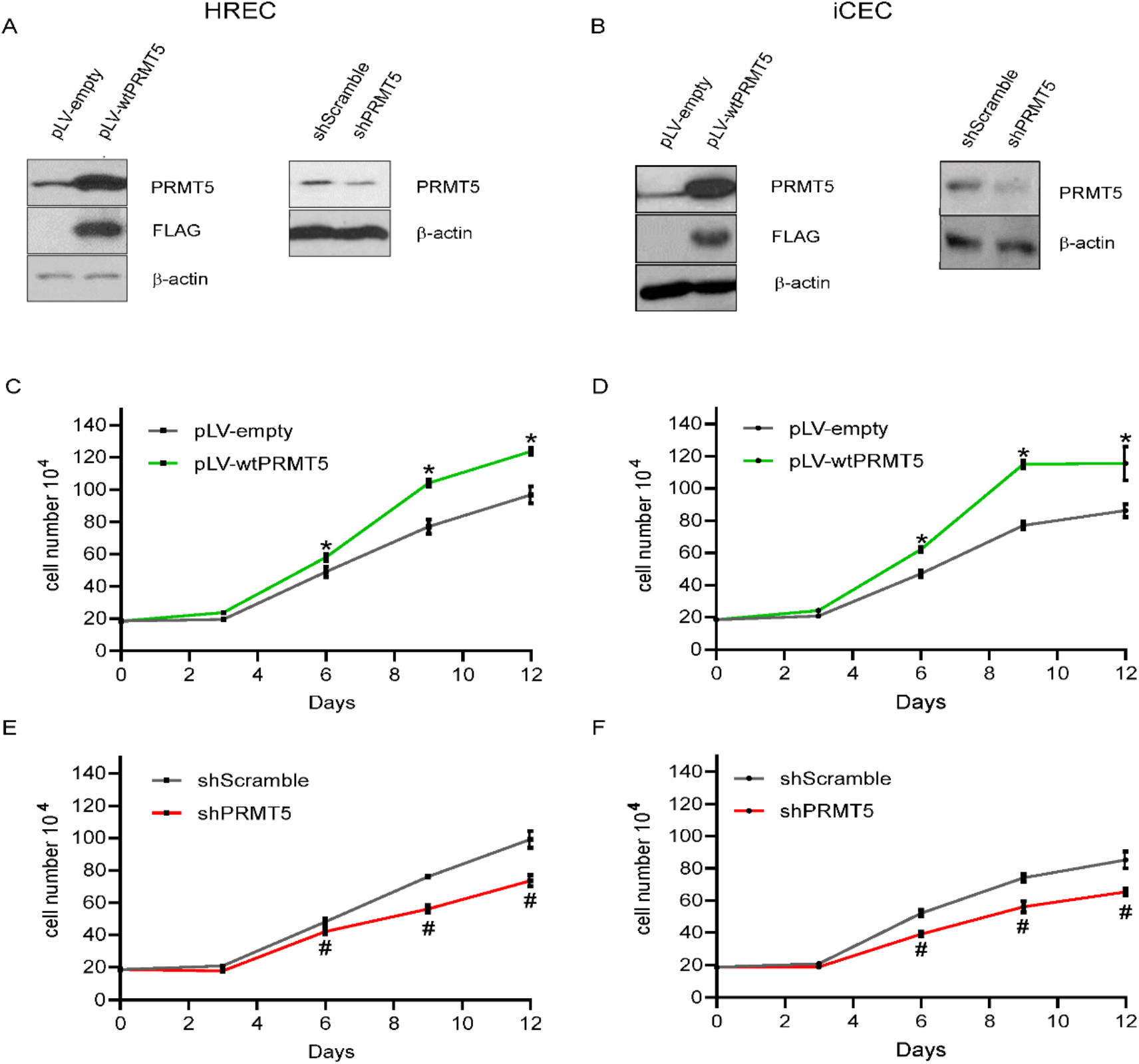
PRMT5 overexpression promotes cell growth. (A) Immunoblot, showing FLAG-tagged wild-type (wt) PRMT5 is successfully overexpressed (*left panel*) or knocked down (*right panel*) in HRECs. (B) Immunoblot, showing FLAG-tagged wtPRMT5 is successfully overexpressed (*left panel*) or knocked down (*right panel*) in iCEC2 cells. (C-F) Effect of PRMT5 on cell proliferation. Overexpression of wtPRMT5 promoted cell growth in HRECs (C) and iCEC2 (D), while shPRMT5 knockdown had the opposite effect in HRECs (E) and iCEC2 cells (F). Mean±SEM, n=3-4 biological replicates. *p<0.05 wtPRMT5 *vs.* pLV-empty; #p<0.05, shPRMT5 *vs.* shScramble, unpaired Student’s *t*-tests with Welch’s correction.

### 3.4 PRMT5 inhibitor PR5-LL-CM01 exhibits antiangiogenic properties in ocular endothelial cells

PR5-LL-CM01 (Figure 4A) has been identified in our lab as an effective PRMT5 inhibitor which downregulates NF-κB activity (Prabhu et al, 2017b). Previously, the effect of PR5-LL-CM01 has been shown only in the context of cancer (Prabhu et al., 2017b). Here, we further studied PR5-LL-CM01 in the context of ocular angiogenesis, in HRECs and iCEC2 cells. Relative to DMSO, PR5-LL-CM01 dose-dependently reduced the proliferation of these cells in a micromolar range, GI_50_ = 2.42 μM in HRECs and 2.98 μM in iCEC2 cells (Figure 4B and C). This antiproliferative effect was evident by the cell cycle arrest in these endothelial cells, where increasing concentrations of PR5-LL-CM01 decreased cells in S phase in comparison to the cells treated with 1% DMSO (Figure 4D-G). Further, a concomitant increase in the cell populations was noted in the G0/G1 phase of the cell cycle on increasing doses of PR5-LL-CM01, suggesting a concentration-dependent inhibition of cell transition from G0/G1 to S-phase.

**Figure 4.**
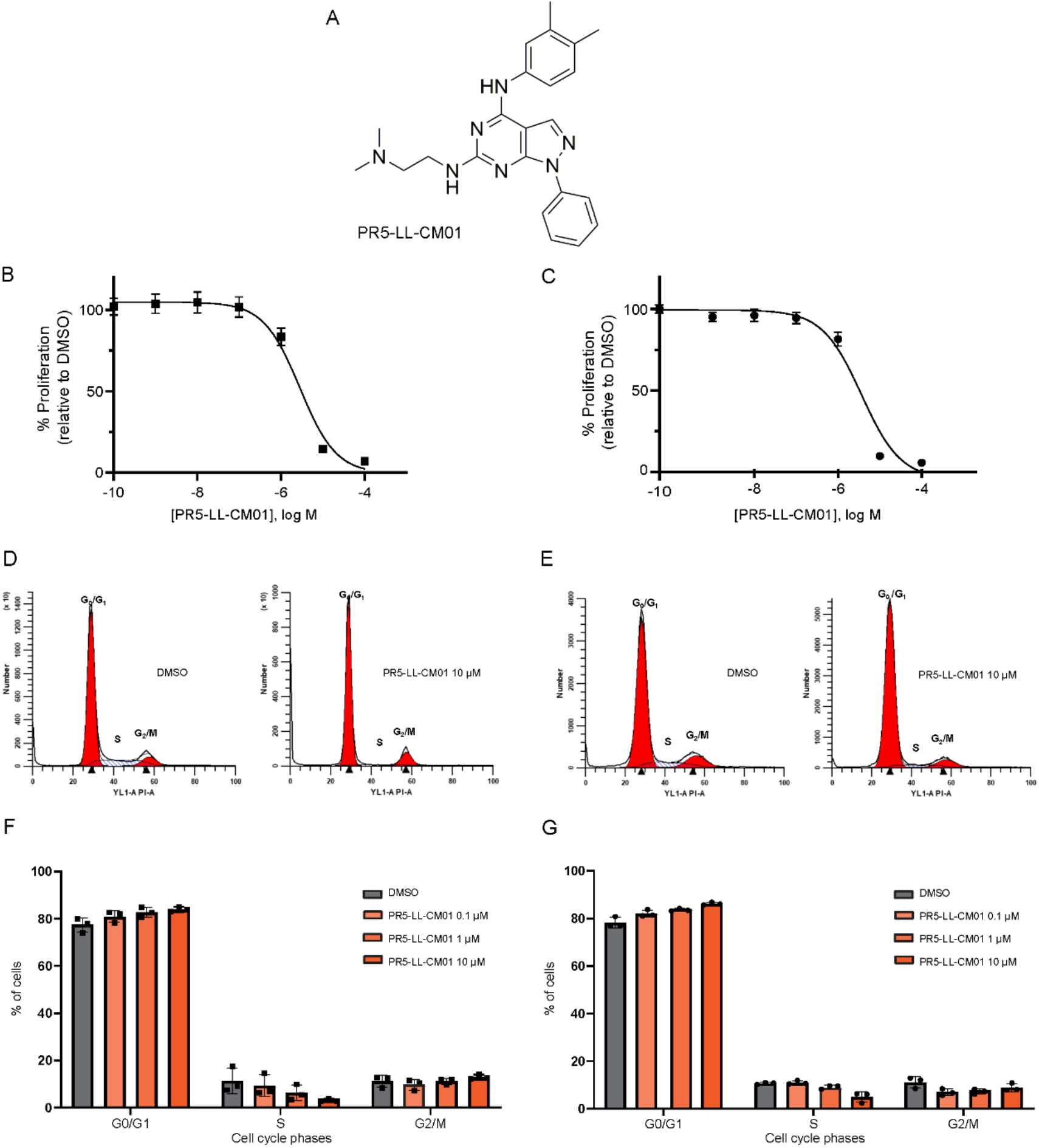
PR5-LL-CM01 is antiangiogenic *in vitro*. (A) Structure of PRMT5 inhibitor PR5-LL-CM01 (Prabhu, et al, 2017b). (B) PR5-LL-CM01 is antiproliferative at 48 hours of treatment on HRECs (B) and on iCEC2 cells (C). Mean±SEM, n=3 technical replicates. Representative data from three biological replicates. PR5-LL-CM01 dose-dependently decreases cells in S-phase and increases cells in G0/G1 after 24 hours of treatment in (D, F) HRECs, and in (E, G) iCEC2 cells. Mean±SEM of percentage of cells, n=3 biological replicates.

### 3.5 PR5-LL-CM01 decreases PRMT5-mediated NF-κB activation and its downstream target gene expression

As PR5-LL-CM01 is known to inhibit NF-κB activity by downregulating PRMT5 activity in cancer (Prabhu et al, 2017b), we sought to determine if PR5-LL-CM01 has a direct effect on the dimethylation of the p65 subunit of NF-κB (p65me2), using p65 overexpressing HRECs and iCEC2 cells. As shown in Figure 5A and B, upon treatment with PR5-LL-CM01, p65me2 was decreased in a dose-dependent manner. This suggests PR5-LL-CM01 inhibited PRMT5-mediated p65me2.

**Figure 5.**
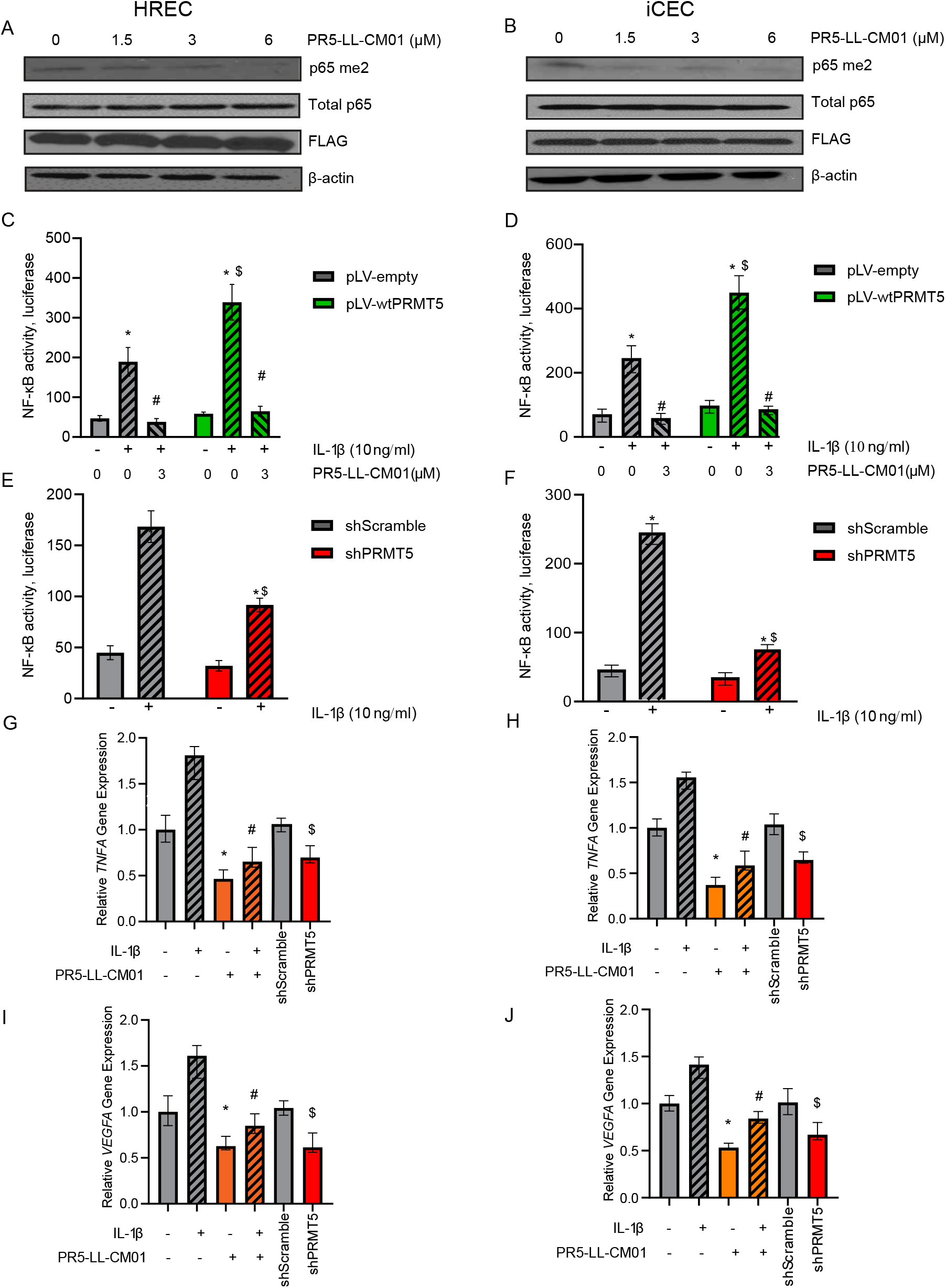
Treatment with PR5-LL-CM01 decreases p65me2, NF-κB activity, and NF-κB target gene expression in HRECs and iCEC2 cells. (A, B) Immunoblots, indicating inhibition of p65-R30me2 by different concentrations of PR5-LL-CM01 treatment in HRECs (A) or iCEC2 cells (B) overexpressing FLAG-tagged wt-p65. (C, D) NF-κB luciferase assay, wtPRMT5 overexpressing HRECs (C) or iCEC2 cells (D) were stimulated with 10 ng/ml IL-1β ± 3 μM PR5-LL-CM01 for 4 hours. PRMT5 overexpression augmented NF-κB induction upon IL-1β treatment, while PR5-LL-CM01 inhibitor treatment reduced this effect. *p<0.05 *vs.* IL-1β untreated group; #p<0.05 vs. IL-1β-induced group; $p<0.05 *vs.* pLV-empty + IL-1β-treated group. (E, F) Both shScramble and shPRMT5 HRECs (E) or iCEC2 cells (F) were stimulated with 10 ng/ml IL-1β for 4 hours. shPRMT5 inhibited NF-κB activity upon IL-1β treatment. *p<0.05 *vs.* IL-1β untreated group; $p<0.05 *vs.* shScramble + IL-1β-treated group. n=3-4 technical replicates. (G, I) HREC vector cells or (H, J) iCEC2 vector cells were stimulated with 10 ng/ml IL-1β ± 3 μM PR5-LL-CM01 for 4 hours. PR5-LL-CM01 significantly decreased the expression of *TNFA* (G, H) and *VEGFA* (I, J) mRNA. *p<0.05 *vs.* IL-1β untreated and PR5-LL-CM01 untreated group; #p<0.05 *vs.* IL-1β-induced group, and PR5-LL-CM01 untreated group; $p<0.05 vs. shScramble group. n=3-4 biological replicates. To test for significant differences, an unpaired Student’s t-test with Welch’s correction was used for comparing two means, and one-way ANOVA with Dunnett’s post hoc tests was used when comparing more than two means.

Given the documented regulation of NF-κB by PRMT5 (Prabhu et al., 2017b), we wished to determine if PRMT5 overexpression or shPRMT5 knockdown altered NF-κB activity *via* a luciferase assay and if this activity could be reduced with the addition of PR5-LL-CM01 in retinal and choroidal endothelial cells. Overexpression of PRMT5 significantly increased IL-1β-induced NF-κB activity, while addition of PR5-LL-CM01 dramatically reduced NF-κB activity in both HRECs (Figure 5C) and iCEC2 cells (Figure 5D). Furthermore, shPRMT5 knockdown decreased IL-1β-induced NF-κB activity compared to respective controls in both HRECs (Figure 5E) and iCEC2 cells (Figure 5F). To further evaluate the effect of PRMT5 inhibition on NF-κB target gene expression, we conducted qPCR analysis of two well-known NF-κB target genes involved in inflammation and angiogenesis, *TNFA* and *VEGFA.* The expression of both *TNFA* (Figure 5G and H) and *VEGFA* (Figure 5I and J) was significantly decreased by either PR5-LL-CM01 treatment or shPRMT5 knockdown as compared to their respective control cells, in both HRECs (Figure 5G and I) and iCEC2 cells (Figure 5H and J).

Taken together, the above data suggest that PR5-LL-CM01 decreases PRMT5-mediated NF-κB activation and its downstream target gene expression, thus presenting promising therapeutic antiangiogenic and anti-inflammatory potential through NF-κB inhibition.

### 3.6 PRMT5 knockdown reduces angiogenic tube formation in ocular endothelial cells

Tube formation is an *in vitro* property of endothelial cells reflective of angiogenic potential *in vivo*. The Matrigel tube formation assay is a widely used, reproducible model system to study either the activation or inhibition of angiogenic pathways in vitro. Knockdown of PRMT5 in various cancer cells can suppress the protumorigenic functions of PRMT5 (Wei et al., 2012; Chiang et al., 2017; Deng et al., 2017; Jiang et al., 2021). Therefore, we sought to assess whether genetic or chemical inhibition of PRMT5 affects tube formation in HRECs and iCEC2 cells. PRMT5 knockdown (Figures 6A-D) in both cell types significantly reduced tube formation ability compared with the shScramble control. Consistent with this, PRMT5 inhibition by PR5-LL-CM01 dose-dependently decreased tube formation in both cell types (Figure 6E-H). These data suggest that inhibition of PRMT5 by PR5-LL-CM01 is comparable with genetic knockdown and both can halt angiogenic properties of cells relevant to ocular neovascular diseases.

**Figure 6.**
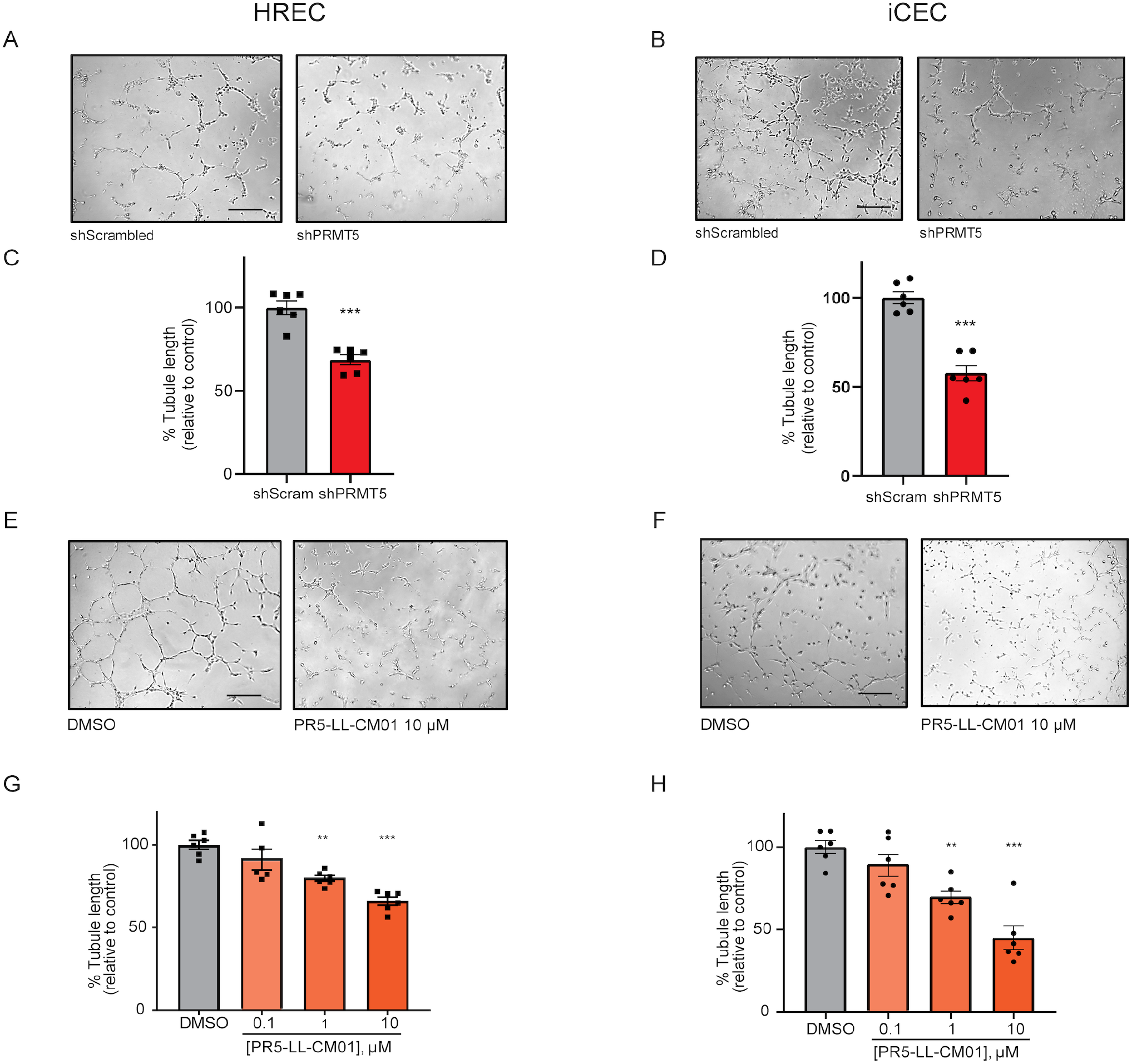
PRMT5 inhibition by shRNA knock down or PR5-LL-CM01 treatment reduces tube formation in HRECs and iCEC2 cells. (A-D) Quantitative analysis of tube formation in HRECs (A, C) or iCEC2 cells (B, D) transduced with shScramble vector or shPRMT5, displaying that knockdown of PRMT5 reduces tube formation ability. ***p<0.0001 *vs.* shScramble control, unpaired Student’s t-test with Welch’s correction. (E-H) Quantitative analysis of tube formation in HRECs (E, G) and in iCEC2 cells (F, H) demonstrates that PR5-LL-CM01 blocks tube formation in a dose-dependent manner. Mean±SEM, n=6 images. **p<0.01; ***p<0.001 *vs.* DMSO control, one-way ANOVA with Dunnett’s post hoc test. Representative data from three biological replicates. Scale bars = 500 μm.

### 3.7 Hypothetical model

Based on the findings from this study, we provide evidence that PRMT5 methylates and activates NF-κB in endothelial cells. This results in the induction of NF-κB downstream genes, known to include cytokines, angiogenesis factors, chemokines, and antiapoptotic genes, whose functions are critical for inflammation and angiogenesis. Thus, using PR5-LL-CM01 to block the activity of PRMT5 has potential to inhibit neovascularization-associated eye diseases (Figure 7).

**Figure 7.**
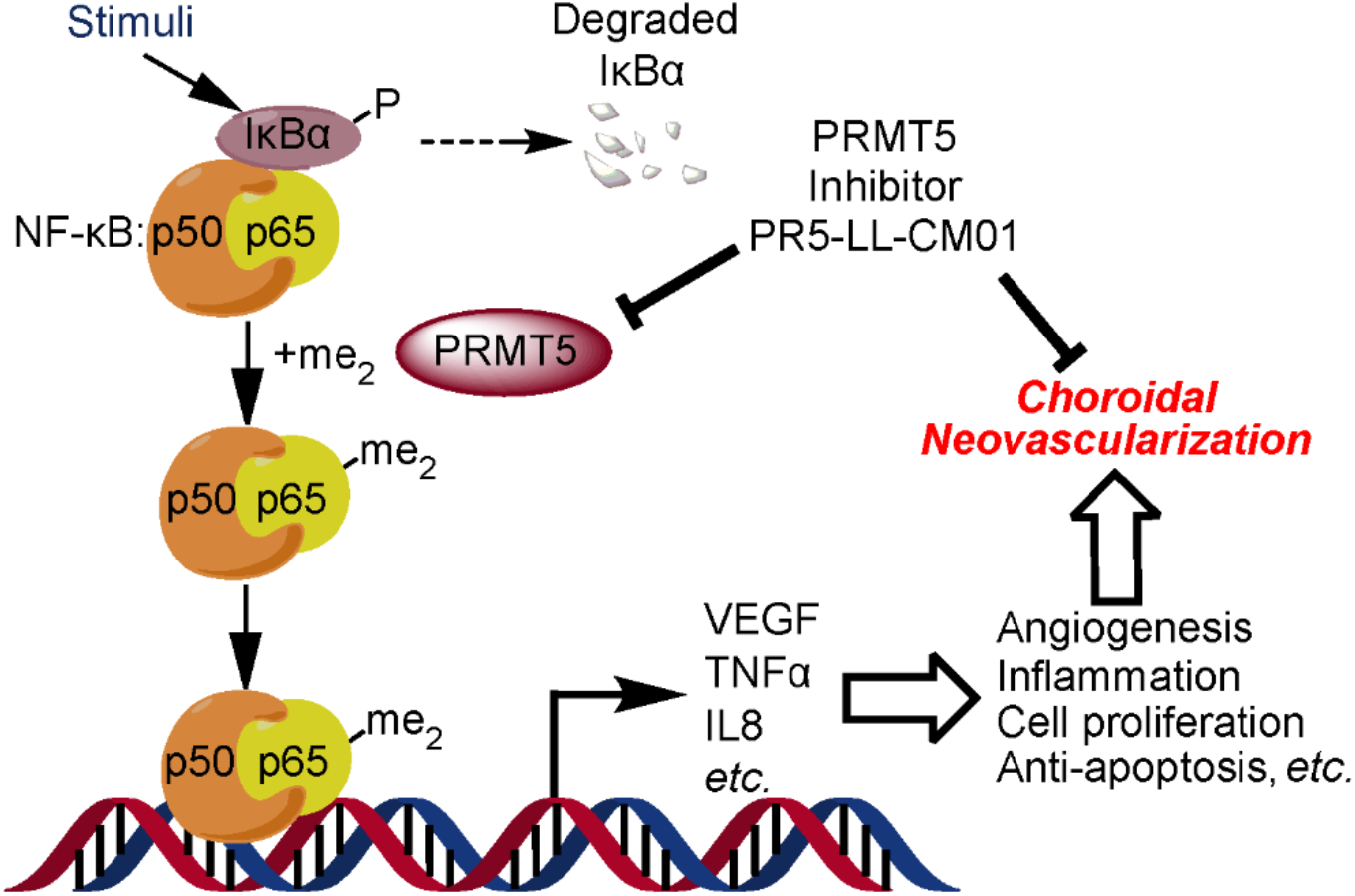
Hypothetical model. PRMT5 methylates and activates NF-κB. This results in the induction of NF-κB downstream genes, including cytokines, angiogenesis factors, chemokines, and antiapoptotic genes, whose functions are critical for inflammation and angiogenesis. Thus, using PR5-LL-CM01 to block the activity of PRMT5 will inhibit neovascularization-associated eye diseases.

## 4 Discussion

Understanding the molecular mechanisms of cellular angiogenic components is crucial for identifying new therapeutics targeting ocular neovascular diseases such as nvAMD, PDR, and ROP. We showed here for the first time that PRMT5 is a novel potential therapeutic target for nvAMD. In this study, we provide experimental evidence that high expression of PRMT5, as observed in the murine L-CNV model and human nvAMD tissues, can impact the regulation of proangiogenic and proinflammatory NF-κB signaling; inhibiting PRMT5 with a potent small molecule inhibitor, PR5-LL-CM01, offers potential therapeutic efficacy.

Our work adds to a body of evidence linking NF-κB signaling with neovascularization. Although the pathophysiology of nvAMD is multifactorial, NF-κB-mediated inflammation and activation of angiogenic VEGF signaling are vital processes that control neovascularization in the ocular system. Growing evidence has shown the influence of NF-κB signaling as a target for nvAMD (Liu et al., 2014; Lu et al., 2014; Ghosh et al., 2017; Hikage et al., 2021). As a master regulator, NF-κB regulates inflammation in the retina under conditions of stress through the transcriptional regulation of various cytokines and angiogenic factors. Such activation of NF-κB inflammatory signaling eventually results in VEGF upregulation (Wang and Hartnett, 2016). Since the endothelial cell microenvironment is the target site of angiogenesis, previous studies have documented the role of increased TNF-α, activation of NF-κB, and higher VEGF levels in angiogenic endothelial cells including human umbilical vein endothelial cells (HUVECs) (Swiatkowska et al., 2005) and human cardiac microvascular endothelial cells (HCMECs) (Chandrasekar et al., 2004). In the context of ocular angiogenesis, the activation of NF-κB and the associated inflammatory responses stimulated by thrombin have been studied in HRECs in the milieu of diabetic retinopathy (Cowan et al., 2014). Studies on human RPE cells have shown increased VEGF levels stimulated either through TNF-α or NF-κB activation (Nagineni et al., 2012; Li et al., 2014; Park et al., 2015; Wang et al., 2016). Wang et al. (2015) intriguingly documented that neo-vessel genesis in choroidal endothelial cells may be caused by TNF-α facilitated NF-κB activation and VEGF production. However, there have not been studies on ocular endothelial cells targeting NF-κB activation triggered by PRMT5-mediated dimethylation.

We began our investigations by assessing the expression of PRMT5 in human AMD and the L-CNV model. This latter model is an extensively used, robust model system wherein the injury induced in Bruch’s membrane and RPE by laser treatment results in neovascularization akin to that observed in the human pathology in terms of position and appearance of choroidal neovessels (Lambert et al., 2013; Shah et al., 2015). L-CNV mouse eyes have high expression of TNF-α, and intravitreal injections of anti-TNF-α antibody treatment reduced the lesion volume induced in conjunction with lowered expression of VEGF in the RPE/choroid (Wang et al., 2016). The role of NF-κB and its activation by TNF-α have been extensively studied in the murine L-CNV model (Shi et al., 2006; Izumi-Nagai et al., 2007; Lu et al., 2014). Furthermore, recent work showed that NF-κB signaling activated microglial cells to drive angiogenesis in mice with L-CNV, and a competitive inhibitor of NF-κB kinase subunit β (IKKβ) diminished the lesion volume through inhibiting NF-κB signaling and VEGF-A expression (Hikage et al., 2021). In L-CNV, we observed higher expression of PRMT5 in retina and choroid, suggesting a potential correlation between PRMT5 and CNV. It is reasonable to speculate that increased PRMT5 expression in L-CNV may enhance NF-κB activation and proinflammatory cytokine production, as we have shown in progression of cancer and metastasis (Wei et al., 2013; Lu and Stark, 2015; Prabhu et al., 2017b). Further molecular evaluation of this hypothesis will be an interesting next step in the future.

Furthermore, PRMT5 is also present throughout the retina and choroid in human eyes and increased in disease, possibly including endothelial and/or microglial cells activated during neovascularization in nvAMD. NF-κB mediates the regulation and function of proinflammatory genes in innate and adaptive immune cells (Liu et al., 2017). Thus, it is possible that the presence of PRMT5 could be correlated with the characteristic increase of the activated microglia and the chorioretinal endothelial cells, as has been implicated before in early and late AMD (Gupta et al., 2003; Zeng et al., 2016). It is worth noting that in murine L-CNV, the expression of PRMT5 was partially co-localized with retinal and choroidal vasculature, suggesting the incidence of proangiogenic events likely instigated by increased VEGF secretion and endothelial cell activation. We will examine detailed cell-type specific expression of PRMT5 in the retina in the future.

PRMT5 can regulate proliferation, differentiation, invasion, and migration of tumor cells (Xiao et al., 2019). Additionally, we have reported that PR5-LL-CM01, a promising inhibitor of PRMT5, has anti-tumor activity *in vitro* and *in vivo* in PDAC and CRC (Prabhu et al., 2017b). In the present study, we showed that inhibition of PRMT5 *via* PR5-LL-CM01 is anti-angiogenic, dose-dependently impeding key angiogenesis properties of proliferation and tubule formation (Yanni et al., 2009; Han et al., 2021) in ocular (retinal and choroidal) endothelial cells. Likewise, PRMT5 blockade by shRNA knockdown reduced proliferation and tube formation in these cells; while overexpression of PRMT5 enhanced proliferation.

A number of earlier studies demonstrated that the activation of NF-κB has a direct link to cell cycle progression (Chen et al., 2001). For instance, NF-κB inhibition impairs the progression of the cell cycle in HeLa cells (Bash et al., 1997; Kaltschmidt et al., 1999) and human glioma cells (Otsuka et al., 1999). Therefore, we speculate that the observed G1/S-phase blockage induced by PR5-LL-CM01 in HRECs and iCEC2 cells might be, at least in part, associated with the PR5-LL-CM01 induced inhibition of PRMT5-mediated NF-κB activation. In the future, it would be interesting to further examine the detailed mechanism of cell cycle arrest mediated by PR5-LL-CM01.

To our knowledge, the mechanistic role of PRMT5 in endothelial cells, notably in ocular endothelial cell biology has never been studied although the dynamic role of PRMT5 in tumorigenesis, NF-κB methylation, and activation has been studied by us (Wei et al., 2013; Prabhu et al., 2017a; Prabhu et al., 2017b) and others (Wang et al., 2018; Kim and Ronai, 2020; Liu et al., 2021). Therefore, we studied PRMT5-regulated NF-κB activity and the expression of its downstream target genes responsible for inflammation and angiogenesis, such as TNF-α and VEGF-A in retinal and choroidal endothelial cells. Using qPCR analysis, we confirmed that both *VEGFA* and *TNFA* genes were downregulated by PR5-LL-CM01 treatment in HRECs and iCEC2 cells. Thus, we demonstrate that inhibition of PRMT5 reduces NF-κB activity and downregulates target genes important for angiogenesis, although the full spectrum of NF-κB target genes regulated by PRMT5 in endothelial cells remains to be explored.

In summary, we have identified a previously unreported role of PRMT5 in ocular angiogenesis. We employed a specific small molecule inhibitor of PRMT5 to inhibit PRMT5-mediated NF-κB activation and angiogenesis *in vitro*, and we compared this effect with evaluation in PRMT5 knocked down ocular endothelial cells. Hence, this study holds a valuable rationale to develop therapies targeting PRMT5-mediated NF-κB activation in treating not only nvAMD but also potentially other ocular neovascular diseases such as PDR and ROP. Future studies could include a thorough molecular investigation of cell-type specific PRMT5 expression and NF-κB regulation, and exploration of other potential transcriptional targets of inflammation and angiogenesis that play a paradigm role in PRMT5 regulation. On the therapeutic side, investigating the effects of PRMT5 inhibition *in vivo* holds promise to develop novel therapeutics targeting PRMT5 for treating ocular neovascular diseases. Interestingly, since we show that PR5-LL-CM01 decreases *VEGFA* gene expression, there may be an avenue for synergistic effects with approved anti-VEGF therapies in the future.

## 6 Author Contributions

Conceptualization: TL, TWC; Methodology: AMu, MM, MS, KS, NRJ, TL, TWC; Resources: AMo, XQ, MEB, LP; Investigation: AMu, MM, MS, KS; Formal analysis: MM, MS, AMu, KS; Visualization: AMu, MM, MS, KS, TWC; Writing – original draft: AMu, MM; Writing – review and editing: AMu, MM, MS, KS, NRJ, AMo, XQ, MEB, LP, TL, TWC; Funding acquisition: TL, TWC.

## 8 Acknowledgments

We thank the Flow Cytometry Core Facility, Indiana University School of Medicine for its support with Modfit software.

## 9 Data Availability Statement

The datasets generated for this study can be obtained from the corresponding authors on reasonable request.

## 11 Figures

